# The Paleozoic assembly of the holocephalian body plan far preceded post-Cretaceous radiations into the ocean depths

**DOI:** 10.1101/2024.08.01.606185

**Authors:** Chase D. Brownstein, Thomas J. Near, Richard P. Dearden

## Abstract

Among cartilaginous fishes, *Holocephali* represents the species-depauperate, morphologically conservative sister to sharks, rays, and skates and the last survivor of a once far greater Paleozoic and Mesozoic diversity. Currently, holocephalian diversity is concentrated in deep-sea species, suggesting this lineage might contain relictual diversity that persisted in the ocean depths. Yet, the relationships of living holocephalians to their extinct relatives and the timescale of diversification of living species remains unclear. Here, we reconstruct the evolutionary history of holocephalians using comprehensive morphological and DNA sequence datasets. Our results suggest that living holocephalians entered and diversified in deep (>1000 m) ocean waters after the Cretaceous-Paleogene mass extinction, contrasting with the hypothesis that this ecosystem has acted as a refugium of ancient cartilaginous fishes. These invasions were decoupled from the evolution of key features of the holocephalian body plan, including crushing dentition, a single frontal clasper, and holostylic jaw suspension, in the Paleozoic Era, and considerably postdated the appearance of the living familes by 150 million years ago during a major period of biotic turnover in oceans termed the Mesozoic Marine Revolution. These results clarify the origins of living holocephalians as the recent diversification of a single surviving clade among numerous Paleozoic lineages.

## Introduction

The *Holocephali*, which contains the chimaeras, elephant sharks, and their relatives, is by far the most species-poor and morphologically conservative of the four major clades of jawed vertebrates [1]. Because of the relative efficiency with which the genomes of living holocephalians-the chimaeras, ghost sharks, and elephant sharks-can be sequenced [2] and because several extinct clades of vertebrates from the Paleozoic with classically recalcitrant relationships [3,4] seem to show affinities to holocephalans [1,5–11], *Holocephali* has provided important clues about the early evolution of jawed vertebrates [1,5,12–15]. A combination of expanding genomic resources [2] and anatomical data from exceptionally preserved fossil members of this clade [1,5–7,15,16] has illuminated the complex early evolution of the characteristic skeletal, soft tissue, and physiological characteristics during the initial radiation of jawed vertebrates.

Holocephalian diversity is currently concentrated in oceanic environments below depths of 200 meters [17–21]. Given their ancient common ancestry with other vertebrates, deepwater chimaeras and ghost sharks lend credence to the hypothesis that mesopelagic and bathypelagic environments have served as a refuge for ancient biodiversity that has become rare or extinct in other ecosystems [22,23]. Evidence from the fossil record [24] and phylogenomic analyses of diverse living deep sea vertebrate clades [25–33] suggest younger origins for these deepwater radiations. Thus, chimaeras and ghost sharks stand as one of the remaining potential relict vertebrates in the bathypelagic and mesopelagic zones.

Despite the importance of holocephalians for outstanding questions in evolutionary biology, the age of living species diversity has only been inferred a handful of times using taxonomically limited samples [34–36] and the relationships of bizarre Mesozoic holocephalian species [37–39] among the living clades and the numerous lineages known from the Paleozoic remains unexplored in the 21^st^ century [3]. This has obscured the origins of the specialized anatomy of living species [15,17] among morphologically disparate extinct holocephalians and the timescale of evolution in these early-diverging jawed vertebrates.

Here, we reconstruct the evolutionary relationships and timescale of divergence of living holocephalians and their fossil records using newly assembled data from the fossil record and DNA sequences from 81% of extant species diversity [40]. Our results establish and contextualize a Jurassic origin of crown *Holocephali* among multiple diversifications of holocephalians throughout the Paleozoic followed by the survival of only two lineages through the end-Triassic mass extinction. Although we find support for a rapid assembly of key features defining the living holocephalian body plan during the Paleozoic evolution of the total clade, we infer with perhaps one exception that all deep-sea chimaeras and ghost shark clades appeared and diversified following the Cretaceous-Paleogene mass extinction 66.02 million years ago [41]. This recent origin for most deepwater holocephalians eliminates the last major contender for a deep sea ‘living fossil’ vertebrate diversity and establishes this environment as a source of both species richness and morphological innovation [20,42] in this lineage.

## Methods

### (a) Systematics

We follow the conventions of the PhyloCode, which was recently formally applied to ray-finned fish systematics [43], in this study for describing clades [44]. In practice, this means that we refer to total clades using the prefix ‘pan-.’ We also follow emerging conventions in italicizing all clade names [43–45].

### (b) Morphological Dataset Construction

In order to test the phylogenetic position of living holocephalians and Mesozoic species among the larger Paleozoic diversity of the total clade, we constructed a new taxon-character matrix by expanding the dataset of Frey et al. [8] using characters from Didier [17] and Patterson [46]. We also edited character states and added 14 living and extinct holocephalian taxa to thoroughly sample holocephalian diversity within and proximal to the crown clade. Newly added data include genus-level scorings for all five living holocephalian genera, as well as for species in the genera †*Acanthorhina*, †*Chimaeropsis*, †*Elasmodectes*, †*Ischyodus*, †*Metopacanthus*, †*Myriacanthus*, and †*Squaloraja*. Due to the lack of available information on their morphology and our focus on incorporating post-Palaeozoic taxa into the analysis, we did not include any representatives of the Paleozoic pan-holocephalian clades †*Eugenodontia* and †*Petalodontiformes*, which mostly include tooth taxa and for which only a handful of holomorphic specimens [10,11,47] of limited phylogenetic informativeness are known due to taphonomic processes. Future discoveries, including the description of unpublished specimens [4], will be needed to resolve the relationships of these Carboniferous and Permian pan-holocephalians; we expect that the phylogenetic matrix employed in this study will contribute to this pursuit. The final matrix included 36 operational taxonomic units coded for 236 characters. Details of sources for character scorings, character additions, removals, and state modifications, and taxon inclusion are included in the Supplementary Information.

### (c) DNA Sequence Dataset Assembly

In order to sample the species diversity of living holocephalians most fully, we targeted sequences for the *cytb*, *COXI*, *ND2*, and *16s* mitochondrial loci on the NCBI repository Genbank. Using this approach, we were able to sample 100% of all living holocephalian genera and species complexes, including all three species in *Callorhinchus*, seven of the nine recognized species in *Rhinochimaeridae*, and 35 of 47 described species of *Chimaeridae* [40], plus two specimens of *Chimaera* sp. with mitogenomes that appear to diverge considerably from other species [48,49]. We concatenated these data into a single set of aligned sequences for subsequent phylogenetic analyses.

### (d) Morphological Phylogenetic Analyses

We analyzed the morphological matrix using a maximum parsimony approach in PAUP* v4.0a [50]. We set the maxillate ‘placoderm’ stem-gnathostome †*Entelognathus primordialis* [51,52] as the outgroup and ran a heuristic search with TBR branch swapping and 1000 addition sequence replicates holding 5 trees from each replicate (Fig. S1). We also analyzed the morphological matrix using a Bayesian approach in MrBayes v.3.2.7a [53], with †*Entelognathus* set as the outgroup. We used an Mk_v_ model with a gamma-distributed rates parameter, ran the analysis for 3.0 × 10^6^ generations, sampled every 1000 generations, and combined the trees with a burn-in of 25% following confirmation of chain convergence using a standard deviation of split frequencies of <0.01 and confirming adequate mixing in Tracer v. 1.7.1 [54]. Trees resulting from the Bayesian analysis were summarized using a 50% majority rule consensus topology.

Next, we conducted a Bayesian tip-dating analysis of our morphological dataset and a revised set of tip dates for included extinct species (see Supplementary Information) under the Fossilized Birth-Death (FBD) Model [55] as implemented in BEAST 2.6.7 [56,57] with the Mk_v_ model of morphological evolution [58]. We again set †*Entelognathus primordialis* as the outgroup and set the origin prior to 443.8 (Ordovician-Silurian boundary), with bounds of 514.1 Ma (median age of crown vertebrates found by two recent genomics papers [59,60]) and 439.0 .0 Ma (the age of the oldest definite crown gnathostome [61]), as no crown vertebrates are known before Cambrian Stage 3 [62] and crown gnathostomes appear to have initially diversified in the middle-late Ordovician based on both genomic [59,60] and morphological [61,63–68] phylogenies. We conducted two independent runs over 1.0 × 10^8^ generations with 1.0 × 10^7^ pre-burnin and checked for convergence of the posteriors and effective sample size values over 200 using Tracer v. 1.7.1 [54]. The resulting posterior tree sets were combined in LogCombiner 2.6.7. with 10% burnin [56] each into a maximum clade credibility tree with median node heights in TreeAnnotator 2.6.6 [56].

### (e) Molecular Phylogenetic Analyses

We conducted maximum likelihood inference on the concatenated sequence alignment using IQ-TREE [69] on the online web server given the size of the dataset [70]. We calculated standard bootstraps over 100 replicates and SH approximate likelihood ratio tests over 1000 replicates to assess support for given nodes. Next, we conducted Bayesian tip-dating phylogenetic analysis of the sequence alignment in BEAST 2.6.6 using the FBD model and including stem-holocephalian and extinct crown holocephalian clades as fossil tip calibrations, with their placements constrained using monophyletic MRCA priors following the results of Bayesian tip-dating analysis of the morphological dataset. We ran the analysis three times independently over 1.0 × 10^8^ generations, each with a 5.0 × 10^7^ pre-burnin, combined posterior tree sets in LogCombiner 2.6.7. after checking for convergence of the posteriors in Tracer 1.7.1 and summarized the trees in a single maximum clade credibility tree with median node heights in TreeAnnotator 2.6.6.

### (f) Ancestral State Reconstructions

Using our tip-dated phylogeny of holocephalians made using the sequence alignment, we conducted ancestral state reconstructions of habitat type and several features of the living holocephalian body plan to understand the timescale of assembly of the morphology of the crown clade and the origin of deep sea lineages in time. We ran all ancestral state reconstructions in the R package phytools [71]. We collected anatomical data from the literature [1,3,8,14,15,17] and personal examination of specimens and data on habitat occupation from the Fishbase database (https://www.fishbase.se/). Habitat levels are based on previous studies examining depth occupation through deep time by animals [31,72]: 0-200 meters was coded as epipelagic, 200-1000 m was coded as mesopelagic, and below 1000 meters was coded as bathypelagic. To accommodate the observation that many living holocephalians occupy multiple regions of the water column, we used a symmetrical polymorphic character model using the polyMk function in phytools. In our ancestral state reconstructions of habitat and key anatomical features, we conducted simulated stochastic mapping over 1000 simulations using the tip-dated phylogeny focusing on living holocephalians and summarized the posterior distribution of reconstructed ancestral states along a single tree.

## Results

### (a) Phylogenetic Analyses of the Morphological Matrix

Our analyses of the 236 character matrix (Figure 1, Figures S1-S2) resolved four major clades in Pan-*Holocephali*, which our Bayesian tip-dated phylogeny estimates diverges from other chondrichthyans 401.41 million years ago (95% HPD: 374.52, 434.44 Ma) in the middle Devonian. Our phylogeny of *Holocephali* largely agrees with previous analyses [8] regarding the relationships of Paleozoic lineages. The relationships we recover for holocephalians are also broadly consistent across the three analytical approaches (parsimony, uncalibrated Bayesian, tip-dated Bayesian analysis). The first holocephalian clade to diverge, †*Symmoriformes*, includes Devonian and Carboniferous taxa [1,8]. Bayesian analyses incorporating (Figure 1) or excluding (Figure S2) fossil age data resolve the monophyly of this clade with weak to moderate posterior support and place the most recent common ancestor (MRCA) of species included in our phylogeny at 371.59 Ma (95% HPD: 359.84, 390.76 Ma). Next to diverge across all analyses are the †*Iniopterygiformes*, an assemblage of marine chondrichthyans with highly unusual body plans that have historically been difficult to place among jawed vertebrates [4,7,9,15]; this clade last shares a MRCA with other holocephalians in the latest Devonian, 364.69 Ma (95% HPD: 341.4, 392.77), in our phylogeny (Figure 1). As in two recent studies [5,8], we infer that †*Kawichthys moodiei* is more closely related to †*Iniopera* than to †*Symmoriiformes* [6].

**Figure 1.**
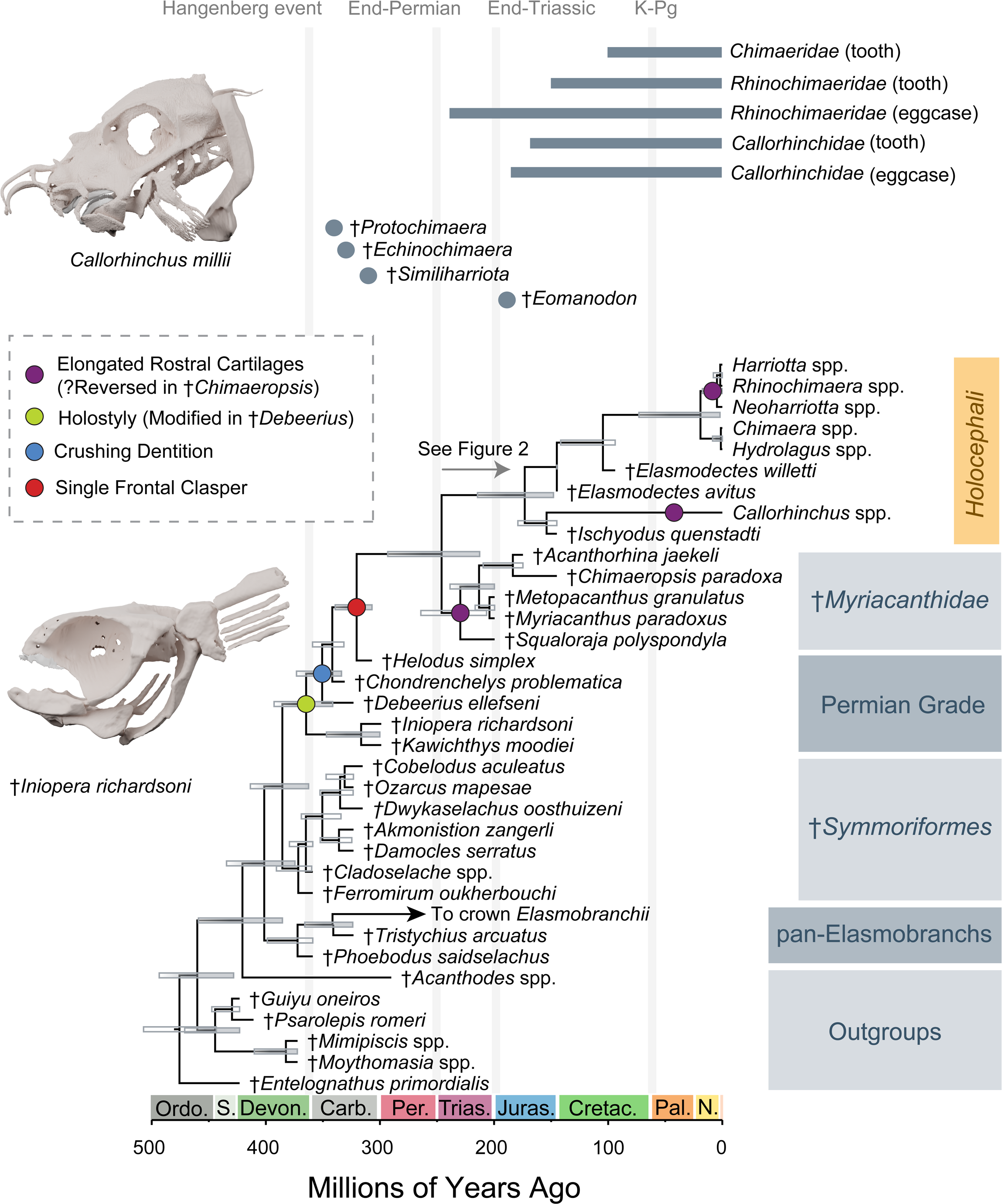
Tip-dated Bayesian phylogeny of living and extinct holocephalians based on morphological characters. Maximum clade credibility tree generated from Bayesian analysis of the morphological matrix and fossil ages in BEAST 2.6.7. Bars at nodes represent 95% highest posterior density intervals for divergence times, and clear bars denote nodes with posterior support values of less than 0.80. Colored dots at nodes indicate inferred origins for key holocephalian features, and CT scan renders spotlight living and extinct holocephalian morphology. Dots and bars above the phylogeny represent the ages of putative crown holocephalians and records of toothplates and eggcases associated with major lineages in the crown group (Supplementary Information).

We resolve the species †*Chondrenchelys problematica* [73] and †*Helodus simplex*, the former of which may represent a larger clade of middle-late Paleozoic species with ethmoid claspers [4,73], in the same positions as found in previous analyses of the Frey et al. [8] dataset. †*Chondrenchelys* is placed closer to the crown clade than †*Debeerius ellefseni*, which appears far more similar to living chimaeras in general body shape [74]. Ancestral state reconstructions of key features of living holocephalians, including the single frontal clasper and durophagy, show that these features first appeared within this grade in the Palaeozoic over a period of approximately 40 million years (Figure 1). †*Helodus simplex* is sister to a long branch extending through the Permian that leads to the two major Mesozoic-Cenozoic holocephalian clades crown *Holocephali* and †*Myriacanthidae*, which we estimate diverged in the Early Triassic 245.97 Ma (95% HPD: 212.66, 239.19 Ma), approximately 5 million years after the End-Permian Mass Extinction. The monophyly of the clade containing crown *Holocephali* and †*Myriacanthidae* is resolved across all analyses with strong bootstrap and posterior support (Figure 1, Figures S1-S2). The crown clade contains species in the living lineages *Callorhinchidae*, *Chimaeridae*, and *Rhinochimaeridae* and the extinct genera †*Ischyodus* and †*Elasmodectes* and is supported with strong bootstrap and posterior support values. The MRCA of the crown clade in our tip-dated Bayesian phylogeny generated using morphological data is placed at 173.14 Ma (95% HPD: 147.94, 214.47 Ma). We resolve the Jurassic species †*Squaloraja polyspondyla*, †*Myriacanthus paradoxus*, †*Metopacanthus granulatus*, †*Acanthorhina jaekeli*, and †*Chimaeropsis paradoxa* as a clade across all analyses with weak support. The oldest available name for this clade is the †*Myriacanthidae* [46].

### (b) Phylogenetic Analyses of the DNA Sequence Dataset

Our analyses of the molecular sequence alignment delimit three major clades of living holocephalians following previous studies [17,34,35]: the elephant sharks in *Callorhinchus*, the longnose chimaeras and spookfishes in *Rhinochimaeridae*, and the chimaeras, rabbitfishes, ratfishes, and ghostsharks in *Chimaeridae* (Figure 2; Figure S3). Although *Chimaeridae* and *Rhinochimaeridae* are found to be sister clades in all analyses (cf. [34,35]), this relationship is weakly supported in both maximum likelihood and Bayesian phylogenies (Figure 2, Figure S3). *Rhinochimaeridae* includes the genera *Harriotta*, *Neoharriotta*, and *Rhinochimaera*, the former of which is found to be paraphyletic with respect to *Rhinochimaera* (Figure 2, Figure S3). *Chimaeridae* includes species traditionally placed in the genera *Chimaera* and *Hydrolagus* (Figure 2; Figure S3). However, as in previous studies we fail to resolve the reciprocal monophyly of these genera (Figure S3) [34,49,75,76]. Instead, we resolve four major clades in *Chimaeridae*, two of which necessitate the resurrection of available generic names.

**Figure 2.**
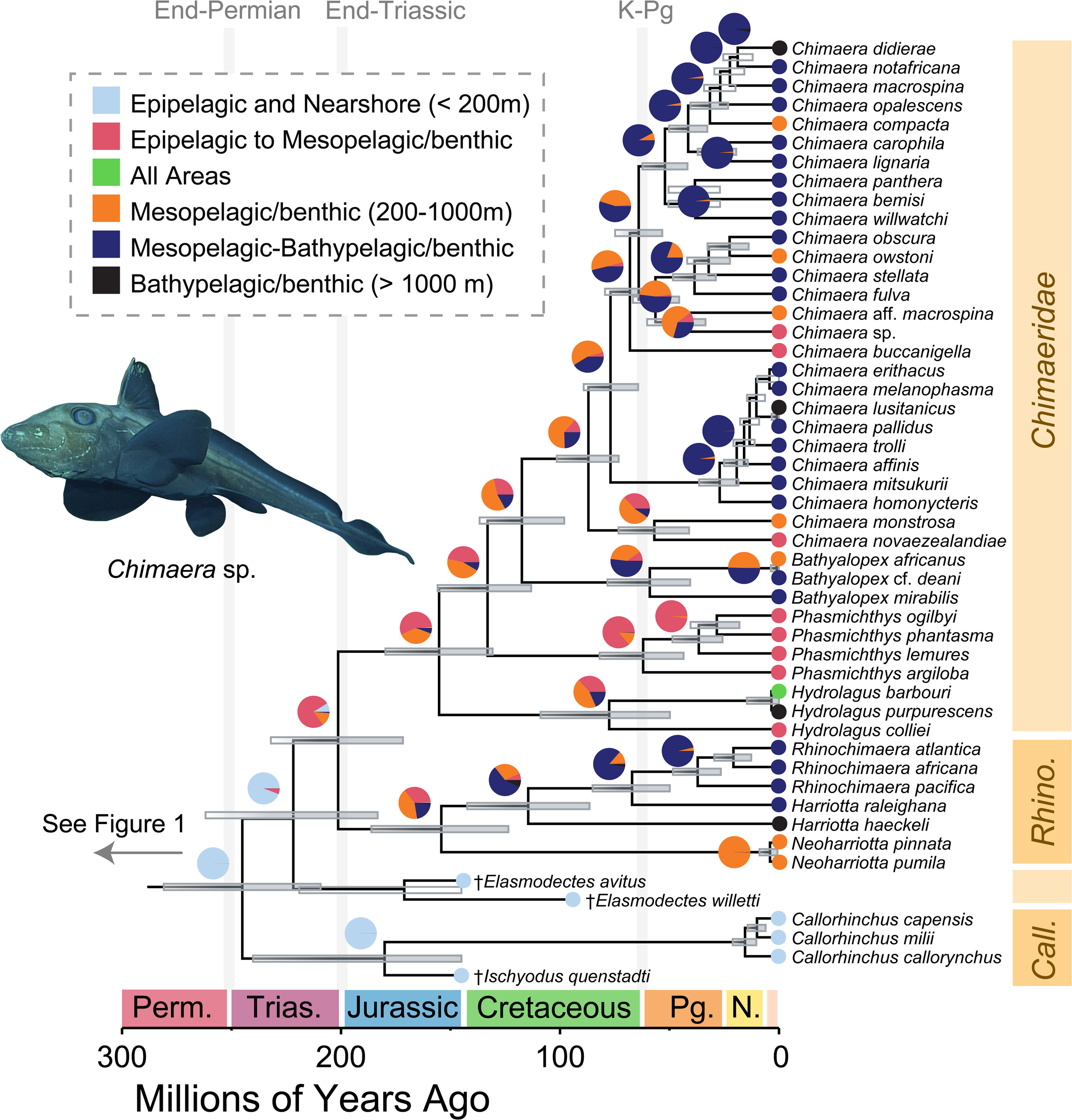
Tip-dated Bayesian phylogeny of living holocephalians. Maximum clade credibility tree generated from Bayesian analysis of the DNA sequence dataset and fixed fossil tip calibrations in BEAST 2.6.7. Bars at nodes represent 95% highest posterior density intervals for divergence times, and clear bars denote nodes with posterior support values of less than 0.80. Pie charts at nodes show reconstructed probabilities for given habitat states, and circles at tips show states associated with each species. The photograph of *Chimaera* sp. is public domain from the NOAA. Abbreviations: *Call*., *Callorhinchus*; *Rhino*, *Rhinochimaeridae*.

The first clade to diverge from other chimaerids is *Hydrolagus*, which includes the type species *H. colliei* and the species *H. barbouri* and *H. purpurescens* (Figure 1, Figure S3). Next to diverge is a clade containing “*Chimaera*” *argiloba*, “*C.*” *phantasma*, “C” *ogilbyi*, and “*H.*” *lemures* (Figure 1, Figure S3). “*Hydrolagus*” *lemures* is the type species of the genus *Phasmichthys* [77], which is the oldest available name for this lineage; we resurrect this genus to include these four species (Figure 1; Figure S3). We also resurrect the genus name *Bathyalopex* [78] for a clade containing “*H. mirabilis*” (the type species of *Bathyalopex*), “*H.*” *africanus*, and “*H.*” *deanii* (Figure 1; Figure S3). This taxonomy restricts the genus *Chimaera* to a clade of at least 25 known species, including the type *C. monstrosa.* All these clades are reciprocally monophyletic with strong support in our maximum likelihood and Bayesian phylogenetic analyses (Figure 1, Figure S3).

We place the divergence of crown holocephalians from their closest relatives in the Early Triassic 245.06 Ma (95% HPD: 209.31, 280.94 Ma) in our tip-dated Bayesian phylogeny made using the DNA sequence dataset. Although it is the earliest diverging living lineage of holocephalians, we place the common ancestor of living species in *Callorhinchus* at 15.49 Ma (95% HPD: 10.31, 21.1 Ma). This result is congruent with the time tree obtained by Inoue et al. [35] and demonstrates that *Callorhinchus* represents the recent diversification of an exceptionally long branch extending from the earliest Mesozoic. In contrast, the clades *Chimaeridae* and *Rhinochimaeridae* last share common ancestry at the Triassic-Jurassic boundary 201.31 Ma (95% HPD: 171.72, 232.05 Ma) in our tip-dated phylogeny. We estimate that both *Chimaeridae* and crown *Rhinochimaeridae* appear in the Kimmeridgian Stage of the Late Jurassic at 155.2 Ma (95% HPD: 130.77, 179.73 Ma) and 154.18 Ma (95% HPD: 123.56, 186.16 Ma), respectively. Except for *Chimaera* and *Hydrolagus*, all chimaerid and rhinochimaerid genera resolved as monophyletic appear after the Cretaceous-Paleogene mass extinction (Figure 2). The major subclades in *Chimaera* also represent post-Cretaceous diversifications (Figure 2).

### (c) Evolution of Habitat Preference

Our ancestral state reconstruction of habitat preference across holocephalian phylogeny broadly finds evidence for multiple recent deep sea invasions and subsequent diversifications rather than an ancient history of life in the meso- and bathypelagic zones. The common ancestor of crown holocephalians is reconstructed as an epipelagic inhabitant with strong support, and it is only after the end-Triassic extinction that mesopelagic clades appear (Figure 2). Except for a clade containing species in the genera *Harriotta* and *Rhinochimaera* with a mid-Cretaceous origin 114.5 Ma, the evolutionary history of bathypelagic habitation dates to after the Cretaceous-Paleogene mass extinction in *Holocephali*. Indeed, our ancestral state reconstruction posits the early Cenozoic as an important period of ecological transition in chimaerid and rhinochimaerid fishes.

Following a transition to partially or fully mesopelagic lifestyles along the backbone of *Chimaeridae*, four major clades transition to the bathypelagic zone. One is *Bathyalopex* (MRCA age: 62.02 Ma). The other three ancestrally bathypelagic clades are species groups in *Chimaera*. The youngest includes the most recent common ancestor of *C. affinis* and *C. homonycteris* and all its descendants (MRCA age: 27.06 Ma), the second youngest includes all species sharing a MRCA with *C. fulva* and *C. obscura* (MRCA age: 38.53 Ma), and the oldest includes all species sharing a MRCA with *C. bemisi* and *C. opalescens* (MRCA age: 52.07 Ma). Notably, there is weak support for an ancestral bathypelagic habitat for the larger clade containing the latter two lineages (Figure 2), which last shares a common ancestor 64.05 Ma (95% HPD: 53.29, 74.84 Ma) and is the sister lineage to *Chimaera buccanigella.* Among other major chimaerid lineages, the ancestral habitat of *Hydrolagus* species is unclear but likely in the mesopelagic and bathypelagic zones, and the ancestral habitat of *Phasmichthys* species is strongly inferred to be the mesopelagic and epipelagic zones (Figure 2).

## Discussion

Here, we have provided the first species-level phylogeny and hypothesis of diversification for the *Holocephali* (Figure 1, Figure 2). Our results highlight that living holocephalian diversity represents one of two clades that emerged from a far more morphologically diverse grade of Paleozoic lineages, including the †*Symmoriformes* and †*Iniopterygiformes*. Together with the †*Myriacanthidae*, which includes extinct genera notable for their elongated rostral cartilages [79] and highly modified cephalic claspers [3,37], our phylogeny shows that crown *Holocephali* originated from a single clade that survived the Permo-Triassic mass extinction. Although there is some evidence from tooth fossils that representatives of †*Symmoriformes* persisted into the Cretaceous [80,81], the identity of these putative Cretaceous symmoriform teeth is controversial [82]. These results highlight the importance of the successive Permo-Triassic and End-Triassic mass extinctions for constraining the phylogenetic and morphological diversity of living holocephalians and their closest relatives.

We resurrect two genera, *Bathyalopex* and *Phasmichthys*, for clades in *Chimaeridae* that diverged from recognized lineages in the Jurassic and Cretaceous (Figure 2). This clarifies the systematics of living holocephalians, particularly confusion surrounding the classification of species traditionally placed in the chimaerid genera *Chimaera* and *Hydrolagus* [75,76]. Our placement of species in the genera †*Ischyodus* and †*Elasmodectes* on the stems of two lineages in crown *Holocephali* with strong nodal support across different analyses also provides a starting point for a reinterpretation of the fossil record of post-Paleozoic holocephalian fishes, which consists almost entirely of isolated tooth plates [83–91], on the basis of skeletal remains.

Previous studies of the phylogeny and timescale of evolution of holocephalians have focused solely on living species [34,35] or Paleozoic fossils [1,5,8] to establish a timescale of chondrichthyan and holocephalian evolution. Our tip-dated Bayesian phylogeny of *Holocephali* infers that the divergence of living holocephalians from their closest relatives, the †*Myriacanthidae*, occurred just after the Permo-Triassic mass extinction. This is consistent with the identification of the Pleinsbachian †*Eomanodon simmsi* as a close relative of crown holocephalians [92]. However, our results call into question the identification of Carboniferous forms such as the holomorphic †*Echinochimaera meltoni* [93], †*Similiharriotta* [94], and the toothplate taxon †*Protochimaera mirabilis* [95] as close relatives of the holocephalan crown-group. Given that our analysis suggests that the similar body shapes of crown holocephalians and Permo-Carboniferous species like †*Debeerius ellefseni* were convergently evolved (Figure 1), it may be that the placement of problematic species like †*Echinochimaera meltoni* close to the crown is affected by homoplastic character evolution.

Our results also place all family- and many genus-level divergences within the holocephalian crown clade and the †*Myriacanthidae* within a period of largescale biotic change called the Mesozoic Marine Revolution (MMR) [96–100]. Although the synchronicity of the Mesozoic diversification of holocephalians with the MMR follows the pattern of diversification in durophagous predator guilds during this event [97,98,100], our ancestral state reconstructions support a Paleozoic origin for the crushing dentition of holocephalians as inferred in previous studies [1,73,74,101] and attested to by a rich fossil record of teeth from the Upper Devonian onwards [102] within an earlier proliferation of durophagous clades [103].

The recent age of living holocephalian species diversity is also notable: 93% of living holocephalian species included in our tip-dated tree diverged from their closest relative in the last 65 million years. This pattern is especially evident in *Callorhinchus*, *Rhinochimaera*, *Neoharriotta*, and the *Chimaera affinis*-*Chimaera pallidus* species group, which all originated between 2 and 27 million years ago (Figure 2). Along with recent analyses of other clades with fossil records reaching back into the Paleozoic, such as lampreys [104,105], our results highlight the recent origins of species diversity in some of the most deeply divergent vertebrate lineages.

The evolutionary history of *Holocephali* presented in this study adds to a growing body of evidence favoring a post-Jurassic origin of deep sea vertebrate diversity. Our integration of fossil and molecular data (Figure 1, Figure 2) supports an slightly older age for crown *Holocephali* than found in previous studies [35], our ancestral state reconstructions show that holocephalians first invaded the bathypelagic zone in the Early Cretaceous, with the majority of transitions into the deep sea occurring after the Cretaceous-Paleogene mass extinction among species in the genus *Chimaera* (Figure 2). These ages are comparable to the Cenozoic ages inferred for some species-rich deep-sea ray-finned fish clades, including anglerfishes [26,106], snailfishes and eelpouts [25,106,107], cods and grenadiers [106,108], and rockfishes [109], and even considerably postdate the ages of deep sea transitions in the teleost clades *Aulopiformes* [28,30,31,110–112], *Stomiiformes* [33], and *Elopomorpha* [30,31,113–115]. Owing to the ancient age of the holocephalian total group, the clades †*Chimaeridae* and †*Rhinochimaeridae* could include some the oldest potential deep sea jawed vertebrate lineages. Consequently, our study eliminates a key vertebrate lineage as a candidate relictual component of deep sea assemblages. Instead, the time-calibrated phylogeny presented in this study posits the Cretaceous-Paleogene mass extinction as a facilitator of chimaerid diversification in the deep sea, perhaps in response to ecological opportunity [116]. Our results also establish that chimaeras likely invaded the bathypelagic zone multiple times, highlighting the complex habitat transitions that have occurred in this species-poor lineage over the last 66 million years of Earth history.

Cartilaginous fishes have been highlighted as a trove of ancient vertebrate diversity [36]. Our results, which suggest that deep sea holocephalian diversity originated relatively recently, show that this habitat, which now faces numerous anthropogenic threats [117–120], has been a cradle of species generation in a lineage containing over 400 million years of unique evolutionary history and with living genera with origins deep in the Mesozoic.

## Acknowledgements and Funding

C.D.B. thanks Joshua Moyer for discussions regarding the evolution of holocephalians and cartilaginous fishes, and Vanessa Rhue for access to the vertebrate paleontology collections of the Yale Peabody Museum. T.J.N. is supported by the Bingham Oceanographic Fund of the Yale Peabody Museum. R.P.D. is supported by funding from the European Union’s Horizon 2020 research and innovation program under the Marie Skłodowska-Curie grant agreement No 101062426.

## Data Availability

All data is available in the Supplementary Information or in the Dryad Repository associated with this manuscript: XXXXXX.

