## Supplementary Text for "The Paleozoic assembly of the holocephalian body plan far preceded post-Cretaceous radiations into the ocean depths"

**Supplementary Text: The rapid assembly of the holocephalian body plan far preceded their deep-sea radiation**

**This file includes:**

Fossil Calibration Justification List

Additional Methods for Assembly of the Morphological Matrix

Supplementary References

Supplementary Figure Captions

**Fossil Calibration Justification List.**

†*Entelognathus primordialis*

**Stratigraphy**:  Xiaoxiang Reservoir, Kuanti Formation, Yunnan Province, China; late Ludfordian Stage of the Silurian [1]. The Ludfordian Stage ranges between 425.6 and 423.0 Ma [2].

**Fossil tip age**: 423.0 Ma.

†*Moythomasia* spp.

**Stratigraphy**: Various, Europe and Australia, all at minimum Frasnian [3], which dates to between 382.7 and 372.2 Ma [2].

**Fossil tip age**: 372.2 Ma.

†*Mimipiscis* spp.

**Stratigraphy**: Gogo Formation, Western Australia [4]; Frasnian, 382.7 to 372.2 Ma [2].

**Fossil tip age**: 372.2 Ma.

†*Psarolepis romeri*

**Stratigraphy**: Xitun Formation, Qujing, East Yunnan, China [5]; Lochkovian, 419.2 to 410.8 Ma [2].

**Fossil tip age**: 410.8 Ma.

†*Guiyu oneiros*

**Stratigraphy**: Qujing, Kuanti Formation, Yunnan Province, China; late Ludfordian Stage of the Silurian [1]. The Ludfordian Stage ranges between 425.6 and 423.0 Ma [2].

**Fossil tip age**: 423.0 Ma.

†*Acanthodes* spp.

**Stratigraphy**: Various units from across the globe; the youngest occurrences are Sakmarian (Early Permian) of Germany [6,7]. The Sakmarian ranges from 293.52 to 290.1 Ma [2].

**Fossil tip age**: 290.1 Ma.

†*Phoebodus saidselachus*

**Stratigraphy**: Madene El Mrakib and Aguelmous, Ibâouane Formation [Lahfira Member, Thylacocephalan Layer][8], southern Maïder region, eastern Anti-Atlas, Morocco; middle Famennian Stage of the Devonian [9], which is between 372.2 and 359.3-358.9 Ma [2,10].

**Fossil tip age**: 358.9 Ma.

†*Tristychius arcuatus*

**Stratigraphy**: Carboniferous Oil Shales, Edinburgh, Scotland; Visean to Surpukhovian Stages of the Carboniferous [11–13], between 346.7 and 323.2 Ma [2].

**Fossil tip age**: 323.2 Ma.

†*Ferromirum oukherbouchi*

**Stratigraphy**: Madene El Mrakib and Aguelmous, Ibâouane Formation [Lahfira Member, Thylacocephalan Layer][8], southern Maïder region, eastern Anti-Atlas, Morocco; middle Famennian Stage of the Devonian [8,9], which is between 372.2 and 359.3-358.9 Ma [2,10].

**Fossil tip age**: 358.9 Ma.

†*Cladoselache* spp.

**Stratigraphy**: Various, mainly known from Cleveland Shale, Ohio, USA, North America; Famennian Stage of the Devonian [6,14], between 372.2 and 359.3-358.9 Ma [2,10].

**Fossil tip age**: 358.9 Ma.

†*Damocles serratus*

**Stratigraphy**: Bear Gulch Formation, central Montana, USA, North America [15]; Surpukhovian Stage of the Carboniferous [14,16–19], 330.9 to 323.2 Ma [2].

**Fossil tip age**: 323.2 Ma.

†*Akmonistion zangerli*

**Stratigraphy**: Bearsden locality, Manse Burn Formation, near Glasgow, Scotland; Surpukhovian Stage of the Carboniferous [14,20], 330.9 to 323.2 Ma [2].

**Fossil tip age**: 323.2 Ma.

†*Dwykaselachus oosthuizeni*

**Stratigraphy**: Base of the Prince Albert Formation, Karoo Supergroup, South Africa; Permo-Carboniferous [21]. We assign an age of 315.2 Ma, which is the base of the Moscovian and approximate midpoint between the Pennsylvanian Epoch of the Carboniferous and end of the Early Permian [2]. However, this fossil could be as young as 280.0 Ma [14].

**Fossil tip age**: 315.2 Ma.

†*Ozarcus mapesae*

**Stratigraphy**:  ARC-07 locality, lower shale member of the Fayetteville Formation, near Leslie, Arkansas, USA; Surpukhovian Stage of the Carboniferous [22], 330.9 to 323.2 Ma [2].

**Fossil tip age**: 323.2 Ma.

†*Cobelodus aculeatus*

**Stratigraphy**: Various Pennsylvanian shales in eastern North America; we use the base of the Moscovian, 315.2 Ma, following previous studies [6,14].

**Fossil tip age**: 315.2 Ma.

†*Kawichthys moodiei*

**Stratigraphy**: 150 km southeast of Kansas City, Missouri, at the boundary of the Haskell Limestone and Robbins Shale Members of the Stranger Formation, Kansas, USA, North America; end Carboniferous, 298.9 Ma at minimum [2,23].

**Fossil tip age**: 298.9 Ma.

†*Iniopera richardsoni*

**Stratigraphy**: 150 km southeast of Kansas City, Missouri, at the boundary of the Haskell Limestone and Robbins Shale Members of the Stranger Formation, Kansas, USA, North America; end Carboniferous, 298.9 Ma at minimum [2,23–25].

**Fossil tip age**: 298.9 Ma.

†*Debeerius ellefseni*

**Stratigraphy**: Bear Gulch Formation, central Montana, USA, North America [17]; Surpukhovian Stage of the Carboniferous [14,16–19], 330.9 to 323.2 Ma [2].

**Fossil tip age**: 323.2 Ma.

†*Chondrenchelys problematica*

**Stratigraphy**: Mumbie Quarry and Glencartholm Fish Beds, Calciferous Limestone, Dumfries and Galloway Region, Scotland; Visean Stage of the Carboniferous [26], 346.7 to 330.9 Ma [2].

**Fossil tip age**: 330.9 Ma.

†*Helodus simplex*

**Stratigraphy**: Fenton, Knowles Limestone, Staffordshire, West Midlands, England [27,28]; near Carboniferous-Permian boundary, 307 Ma [2,14].

**Fossil tip age**: 307.0 Ma.

†*Squaloraja polyspondyla*

**Stratigraphy**: Lower Lias, Dorset, England and Osteno lens, Moltrasio Limestone Formation, Como, Italy; lower Sinemurian [29], 199.5 Ma [2].

**Fossil tip age**: 199.5 Ma.

†*Myriacanthus paradoxus*

**Stratigraphy**: Lower Lias, Dorset, England [30]; lower Sinemurian [29], 199.5 Ma [2].

**Fossil tip age**: 199.5 Ma.

†*Metopacanthus granulatus*

**Stratigraphy**: Lower Lias, Dorset, England [30]; lower Sinemurian [29], 199.5 Ma [2].

**Fossil tip age**: 199.5 Ma.

†*Chimaeropsis paradoxa*

**Stratigraphy**: Various localities, Solnhofen Plattenkalk, Altmühltal Formation, Solnhofen, Germany; Tithonian [31], 149.2 to 145.0 Ma [2].

**Fossil tip age**: 145.0 Ma.

†*Acanthorhina jaekeli*

**Stratigraphy**: Holzmaden, Poseidonia Shale, Germany; Toarcian [30,32], 184.2 to 174.7 Ma [2].

**Fossil tip age**: 174.7 Ma.

†*Ischyodus quenstadti*

**Stratigraphy**: Various localities, Solnhofen Plattenkalk, Altmühltal Formation, Solnhofen, Germany; Tithonian [31], 149.2 to 145.0 Ma [2].

**Fossil tip age**: 145.0 Ma.

†*Elasmodectes avitus*

**Stratigraphy**: Various localities, Solnhofen Plattenkalk, Altmühltal Formation, Solnhofen, Germany; Tithonian [31], 149.2 to 145.0 Ma [2].

**Fossil tip age**: 145.0 Ma.

†*Elasmodectes willetti*

**Stratigraphy**: Various localities, English Clay, England and Ain El Kerma, Jebel Tselfat, near Fez, Morocco; Cenomanian [33–35], 100.5 to 93.9 Ma [2].

**Fossil tip age**: 93.9 Ma.

**Methods for assembly of morphological matrix.**

The morphological matrix was built by adapting that of Frey et al. [8] and adding 14 additional taxa and new characters compiled from [30,36] and additional observations from the literature. A full list of character changes and additional characters is given in the supplementary materials.

The following taxa were removed: †*Youngolepis,* †*Cheirolepis,* †*Raynerius,* †*Latviacanthus,* †*Climatius,* †*Brachyacanthus,* †*Parexus,* †*Ptomacanthus,* †*Brochoadmones,* †*Kathemacanthus,* †*Lupopsyrus,* †*Obtusacanthus,* †*Gyracanthides,* †*Tetanopsyrus,* †*Culmacanthus,* †*Uraniacanthus,* †*Diplacanthus,* †*Rhadinacanthus,* †*Ischnacanthus,* †*Cheiracanthus,* †*Halimacanthodes,* †*Mesacanthus,* †*Homalacanthus,* †*Gladbachus,* †*Gydoselache,* †*Pucapampella,* †*Doliodus,* †*“Gutterensis”,* †*Cladodoides,* †*Tamiobatis,* †*Diplodoselache,* †*Orthacanthus,* †*Triodus,* †*Thrinacodus,* †*Gogoselachus,* †*Acronemus,* †*Homalodontus,* †*Synechodus,* †*Egertonodus,* †*Hamiltonichthys,* †*Tribodus,* †*Onychoselache*, *Chimaeroidei*

The following taxa were added: *Callorhincus, Chimaera, Hydrolagus, Rhinochimaera, Harriotta, Neoharriotta,* †*Isychodus quenstadti,* †*Elasmodectes avitus,* †*Elasmodectes willetti,* †*Chimaeropsis paradoxus,* †*Acanthorhina,* †*Metopacanthus,* †*Myriacanthus,* †*Squaloraja*

**Sources for morphological character scoring**

| **Taxon** | **Reference** |
| --- | --- |
| †*Entelognathus* | [1,37] |
| †*Mimipiscis* | [38] |
| †*Moythomasia* | [38] |
| †*Guiyu* | [39–42] |
| †*Psarolepis* | [5,43,44] |
| †*Acanthodes* | [7,45–47] |
| *Squalus* | Pers. Obs. (CDB), FLMNH ICH 34948 (via Morphosource) [48,49] |
| †*Phoebodus* | [9] |
| †*Tristychius* | [12,13,50] |
| †*Akmonistion* | [20] |
| *Callorhincus* | [36,51,52] |
| *Chimaera* | [36] |
| *Hydrolagus* | [36] |
| *Rhinochimaera* | [36] |
| *Harriotta* | [36] |
| *Neoharriotta* | [36] |
| †*Ischyodus quenstadti* | [31,53,54] |
| †*Elasmodectes avitus* | [31,53,54] |
| †*Elasmodectes willetti* | [33,34] |
| †*Chimaeropsis paradoxus* | [30,31,53] |
| †*Acanthorhina* | [30,32] |
| †*Metopacanthus* | [30] |
| †*Myriacanthus* | [30] |
| †*Squaloraja* | [29,30] |
| †*Chondenchelys* | [26,27] |
| †*Cladoselache* | Pers. Obvs (CDB) [55–58] |
| †*Cobelodus* | [59,60] |
| †*Damocles* | [12,13] |
| †*Debeerius* | [17] |
| †*Dwykaselachus* | [14] |
| †*Ferromirum* | [8] |
| †*Helodus* | [27,30,61] |
| †*Iniopera* | [24,52,62,63] |
| †*Kawichthys* | [23] |
| †*Ozarcus* | [22,59] |

### Morphological Characters

The following characters removed, as they were uninformative for the reduced taxon list (all one state). Apomorphic characters, while uninformative, were retained. Characters with numbers from Frey et al. [9]: 29, 33, 45, 75, 77, 93, 108, 112, 128, 138, 140, 149, 161, 186, 190, 192, 194, 212, 217, 222, 223, 224, 226, 228, 230

The following characters were added or modified: 24, 25, 26, 39, 40, 41, 53, 54, 79, 80, 81, 82, 83, 84, 85, 108, 109, 110, 114, 126, 161, 213, 215, 222, 225, 231,

Added characters and modified characters are written in bold text. Additionally added characters are marked with an asterisk.

Abbreviations for character sources:

Did95, Didier 1995

Mai86, Maisey 1986

Pat65, Patterson 1965

*Skeletal tissues*

1. Tessellate calcified cartilage: absent (0); present (1).
2. Perichondral bone: present (0); absent (1).
3. Extensive endochondral ossification: absent (0); present (1).
4. Extensive calcified cartilage: absent (0); present (1).
5. Tubular dentine: absent (0); present (1).
6. Pore canal network: absent (0); present (1).
7. Acrodin tooth caps (enameloid cap restricted to crown apex): absent (0); present (1).

*Squamation & related structures*

8. Trunk scales monocuspid (0); multicuspid (1).
9. Scale growth concentric: absent (0); present (1).
10. Peg-and-socket articulation: absent (0); present (1).
11. Anterodorsal process on scale: absent (0); present (1).
12. Body scales with bulging base: absent (0); present (1).
13. Body scales with flattened base: absent (0); present (1).
14. Body scales with basal canal or open basal vascular cavity: absent (0); present (1)
15. Neck canal: absent (0) present (1).
16. Cranial sensory line canal passes between or beneath scales (0); passes over scales and/or is partially enclosed or surrounded by scales (1); perforates and passes through scales (2).
17. Postcranial sensory line canal passes between or beneath scales (0); passes over scales and/or is partially enclosed or surrounded by scales (1); perforates and passes through scales (2).
18. Lepidotrichia: absent (0); present (1).
19. Fringing fulcra: absent (0); present (1).
20. Scute-like ridge scales (fulcra): absent (0); present (1).

*Cranial dermal skeleton*

21. Cranial cap denticles, single-crowned, non-growing: absent (0); present (1).
22. Sclerotic ring: absent (0); present (1).
23. Number of sclerotic plates: four or less (0); more than four (1).
24. **Dermal skull roof includes large dermal plates (0); consists of plates, tesserae or scales (1)**

Character state 2 has been removed and replaced with the subsequent character to allow for the fact that in taxa with a naked skull roof the squamation is micromeric (i.e. reduced to lateral line scales). The cranial covering in †*Acanthorhina* is no longer present and may have been removed during preparation (Duffin, 1983): it is scored unknown (?).

25. ***Micromeric skull roof reduced: absent (0); present (1)**
26. ***Elements of micromeric skull roof consolidated into plates: absent (0); present (1)**

In several Mesozoic holocephalians the scales of the cranial roof are consolidated into tuberculated plates [30]. In †*Squaloraja*, the head is more heavily covered in denticles than in extant chimaeroids but these are not consolidated into plates: it is scored 0 [29,30]

27. Dermal ornamentation: smooth (0); parallel, vermiform ridges (1); concentric ridges (2); tuberculate (3).
28. Cranial tessera morphology: large interlocking plates (0); microsquamose, no larger than body squamation (1).

Extinct *Holocephali* with enlarged scales on the skull roof are coded as state 0

29. Anterior or mesial edge of nasal notched for anterior nostril: absent (0); present (1).
30. Supraorbital: absent (0); present (1). [1,39].
31. Large median bone contributes to posterior margin of skull roof: absent (0); present (1).
32. Pineal opening perforates dermal skull roof: present (0); absent (1).
33. Consolidated cheek plates: absent (0); present (1).
34. Dermal intracranial joint: absent (0); present (1).
35. Sensory line network preserved as open grooves (sulci) in dermal bones (0); sensory lines pass through canals enclosed within dermal bones (1).
36. Sensory canal or pit-line associated with maxilla: absent (0); present (1).
37. Jugal portion of infraorbital canal joins supramaxillary canal: present (0); absent (1).
38. Anterior pit line of skull roof: absent (0); present (1).
39. ***Supraorbital sensory line meets supratemporal commissure caudally: absent (0); present (1)**

In most gnathostomes the caudal connection of the supraorbital line is to the infraorbital line. In extant chimaeras it instead connects to the supratemporal commissure [36,64]. This connection is also present in †*Squaloraja* [29] and in †*Ischyodus* [54]. A reconstruction of the cranial sensory lines in †*Debeerius* appears to show this same anatomy [17] and it is scored here as present.

40. ***Angular lateral line: absent (0); present (1)**

In most gnathostomes a single lateral line, the infraorbital line, lies between the orbit and the mouth. In holocephalians a second line, the angular lateral line, is present ventral to the infraorbital line [36,64]. This line is also present in †*Squaloraja* [29,30] and is tentatively reconstructed in †*Isychodus* [54]. A reconstruction of the cranial sensory lines in †*Debeerius* lacks a second canal below the orbit and it is scored here as absent.

41. ***Lateral line canals on rostrum enlarged with expanded dilations: absent (0); present (1)**

Did95 (Ch98)

This is a feature of chimaerid lateral lines [36]. It appears to be absent in †*Squaloraja* [29,30].

42. Spiracular opening in dermal skull roof bounded by bones carrying otic canal: absent (0); present (1).
43. Dermohyal (submarginal) ossification: absent (0); present (1).
44. Branchiostegal series: absent (0); present (1).
45. Opercular and subopercular bones: absent (0); present (1).
46. Branchiostegal plate series along ventral margin of lower jaw: absent (0); present (1).
47. Branchiostegal ossifications plate-like (0); narrow and ribbon-like (1); filamentous (2).
48. Branchiostegals imbricated: absent (0); present (1).
49. Opercular cover of branchial chamber complete or partial (0); separate gill covers and gill slits (1).
50. Gular plates: absent (0); present (1).

*Hyoid and gill arches*

51. Gill skeleton mostly beneath otico-occipital region (0); mostly posterior to occipital region (1).
52. First branchial arch meets neurocranium ventral to otic region (0); posterior to otic region (1)
53. ***Hyoid arch articulates with neurocranium: absent (0); present (1)**

Absent in extant holocephalians

54. ***Pharyngohyal: absent (0); present (1)**

Did95 (Chs 54)

A pharyngohyal is present in extant holocephalians and has also been described in †*Debeerius* [17].

55. Perforate hyomandibula: absent (0); present (1).
56. Interhyal: absent (0); present (1).
57. Ceratohyal with posterior/proximal external fossa: absent (0); present (1).
58. Ceratohyal with broad posteroventral flange or shelf, projecting laterally into recess in Meckel's cartilage: absent (0); present (1).
59. Ceratohyal spatulate or bladed anteriorly: absent (0); present (1).
60. Hypohyals: absent (0); present (1).
61. Basihyal: absent, hyoid arch articulates directly with basibranchial (0); present (1).
62. Separate supra- and infra-pharyngobranchials absent (0); present (1).
63. Pharyngobranchials directed anteriorly (0); posteriorly (1).
64. **Pharyngobranchials III-V fused into single complex unit: absent (0); present (1)**

Modified from Did95 (Ch. 56)

In extant holocephalians the posterior three pharyngobranchials are fused into a complex element with the fourth and fifth epibranchials. This appears to be absent in †*Debeerius* [17].

65. Posteriormost branchial arch bears epibranchial unit: absent (0); present (1). Scored as uncertain for †*Ferromirum* [8].
66. Epibranchials bear posterior flange: absent (0); present (1).
67. Hypobranchials directed anteriorly (0); hypobranchials of second and more posterior	gill arches directed posteriorly (1).
68. Multiple unpaired basibranchial mineralisations absent (0); present (1).
69. Elongate posterior copula projects posteriorly, beyond rearmost branchial arch: absent (0); present (1).

*Dentition & tooth-bearing bones*

70. Oral dermal tubercles borne on jaw cartilages: absent (0); present (1).
71. Pharyngeal teeth or denticles: absent (0); present (1).
72. Tooth families/generative tooth sets: absent (0); present (1).
73. Tooth families/generative sets restricted to symphysial region (0); distributed along jaw margin (1).
74. Number of generative tooth sets per jaw ramus: 15 or fewer (0); 20 or more (1).
75. Bases of tooth families/generative sets: single, continuous plate (0); some or all whorls consist of separate tooth units (1).
76. Lingual torus: absent (0); present (1).
77. Basolabial shelf: absent (0); present (1).
78. Teeth with three slim main cusps almost equal to each other, strongly recurved: absent (0); present (1).
79. Toothplates absent (0); present (1).
80. **Paired toothplate complement restricted to two pairs in the upper jaw and a single pair in the lower jaw: absent (0); present (1).**

This character has had the word “paired” added to it. This is variable in myriacanthoids: in †*Metopacanthus* there are three toothplates in the upper jaw and it is scored 0. We follow Patterson’s interpretation of the toothplate complement in †*Acanthorhina* and score it and †*Chimaeropsis paradoxa* present 1 [30,31,65].

81. ***Hypermineralised regions on all toothplates (tritors) present on all toothplates: absent (0); present (1)**

Did95 (Chs 52)

In chimaeroid teeth hypermineralised tissue is organised into distinct regions (tritors) across the surface of all toothplates [36,66]. In some Mesozoic forms these are absent (†*Squaloraja*) or very simple and restricted only a subset of toothplates (other myriacanthids)[30]. However, they are present in †*Ischyodus* and in *Elasmodectes* [67]. Only applicable to taxa with toothplates.

82. ***Tritoral dentine organised into rod-like structures: absent (0); present (1)**

Did95 (Ch. 89)

These structures, ‘tritoral rods’, are present in chimaerids but not in *Callorhincus* [36]. Among extinct taxa in our matrix, they are present in †*Elasmodectes*. This is only applicable to taxa with tritors on all toothplates [67].

83. ***Toothplates have prominent descending lamina: absent or reduced (0); present (1)**

Did95 (Ch. 53)

A prominent descending lamina is present in the toothplates of myriacanthids, †*Ischyodus*, and *Callorhinchus* [36,68]. In chimaerids, this lamina is weakly developed and not present in all toothplates.

84. ***Vomerine toothplates with several rows of parallel ridges exposed on posterior face of occlusal surface: absent (0); present (1)**

Did95 (Ch. 103)

This character unites *Chimaeridae* [36].

85. ***Symphyseal toothplate: absent (0); present (1)**

Pat65

Myriacanthids have a symphyseal, chisel-like toothplate on the lower jaw [30]. This is also present in †*Acanthorhina* and in †*Chimaeropsis* [31,65,68].

86. Dermal plates on biting surface of jaw cartilages: absent (0); present (1).
87. Gnathal plates mesial to and/or above (or below) jaw cartilage: absent (0); present (1).
88. Maxilla and premaxilla *sensu stricto* (upper gnathal plates lateral to jaw cartilage without palatal lamina): absent (0); present (1).
89. Dentary bone encloses mandibular sensory canal: absent (0); present (1).
90. Infradentary foramen and groove, series: absent (0); present (1).
91. Tooth-bearing median rostral: absent (0); present (1).
92. Median dermal bone of palate (parasphenoid): absent (0); present (1).
93. Denticulated field of parasphenoid: without spiracular groove (0); with spiracular groove (1).
94. Denticle field of parasphenoid with multifid anterior margin: absent (0); present (1).

*Mandibular arch*

95. Large otic process of the palatoquadrate: absent (0); present (1).
96. Oblique ridge or groove along medial face of palatoquadrate: absent (0); present (1).
97. Fenestration of palatoquadrate at basipterygoid articulation: absent (0); present (1).
98. Perforate or fenestrate anterodorsal (metapterygoid) portion of palatoquadrate: absent (0); present (1).
99. Articulation surface of the palatoquadrate with the postorbital process directed anteriorly (0); laterally (1); dorsally (2).
100. Palatoquadrate fused to the neurocranium: absent (0); present (1).
101. Mandibular knob or mesial process: absent (0); present (1).
102. Jaw articulation located on rearmost extremity of mandible: absent (0); present (1).
103. Meckel's cartilage with flange or shelf projecting posteriorly from the lateral cotylus (glenoid): absent (0); present (1).
104. Dental trough adjacent to oral rim on Meckel’s cartilage and palatoquadrate: absent (0); present (1).
105. Dental trough divided, scalloped tooth-bearing margin on Meckel’s cartilage and palatoquadrate: absent (0); present (1).
106. Mandibular symphysis fused: absent (0); present (1).
107. ***Plate or spine on posterior end of mandible: absent (0); present (1)**

Pat65

†*Myriacanthus* and †*Metopacanthus* have a distinctive dermal spine on the posterior part of the Meckel’s cartilage [30]. †*Acanthorhina* has a dermal plate in a similar position and we score this present 1 This appears to be absent in †*Squaloraja*. In †*Chimaeropsis* a dermal plate associated with the back of the mouth may be a mandibular spine but this is unclear: it is scored unknown ?.

*Neurocranium*

108. ***Rostral cartilage: reduced or absent (0); supports plough-shaped snout (1) elongate supporting long anterior process (2)**

Did95 (Chs 77, 92, 99)

This character captures the shape of the rostrum in callorhinchids and rhinochimaerids respectively. It is scored inapplicable for taxa with dermal plates covering the snout. †*Ischyodus* was interpreted by as [54] as having a shark-like snout, however, the shape approximates that of *Callorhinchus*: we have scored it as state 1. In †*Squaloraja*, †*Metopacanthus*, and †*Acanthorhina* the snout is elongate, and it is unclear how much of this is rostral cartilage vs ethmoid braincase: nonetheless these are scored as state 2. For †*Myriacanthus* we follow Pattersons’s interpretation of NHM PV P 10130 as having had an elongate rostral cartilage and frontal clasper [30].

109. ***Frontal tenaculum: absent (0); present (1)**

Did95 (Ch. 63)

A frontal tenaculum is present in extant holocephalians and is also present in †*Helodus* We follow the interpretation of the cartilage over the ethmoid region of *Acanthorhina* as being the base of a frontal tenaculum [65]

110. ***Frontal tenaculum: small cartilage (0); enlarged to extend to rostral tip (1)**

In †*Metopacanthus* and †*Squaloraja*, the frontal tenaculum is massively enlarged, extending along the rostrum. †*Isychodus* is scored 0 for this character, although the frontal tenaculum is reconstructed as larger than in living holocephalians [54].

111. Internasal vacuities: absent (0); present (1).
112. Precerebral fontanelle: absent or minimal (0); present and large (1).
113. Space for forebrain and (at least) proximal portion of olfactory tracts narrow and elongate, extending between orbits: absent (0); present (1).
114. ***Ophthalmic nerves carried from orbit to rostrum by canal/s: absent (0); present (1)**

Modified from Patterson (Ch. 2)

In elasmobranchs the anterior route of the ophthalmic nerves passes out of the anterodorsal wall of the orbit and onto the cranial roof. In holocephalians they instead travel through an ethmoid canal: an unpaired passage that conveys the ophthalmic nerves through the ethmoid region before they exit through the rostral foramen [26,30]. Paired canals interpreted as serving the same function this have been identified in the Palaeozoic holocephalian †*Helodus* [61].

115. Rostral bar: absent (0); present (1).
116. Internasal groove absent (0); present (1).
117. Orbitonasal lamina expanded: absent (0); present (1).
118. Elongate, tooth-bearing, pre-nasal ethmo-rostral region: absent (0); present (1).
119. Palatobasal (or orbital) articulation posterior to the optic foramen (0); anterior to the optic foramen, grooved, and overlapped by process or flange of palatoquadrate (1); anterior to optic foramen, smooth, and overlaps or flanks articular surface on palatoquadrate (2).
120. Supraorbital shelf broad with convex lateral margin: absent (0); present (1).
121. Interorbital space broad (0); narrow (1).
122. Optic pedicel: absent (0); present (1).
123. Ophthalmic foramen in anterodorsal extremity of orbit communicates with enclosed cranial space: absent (0); present (1).
124. Extended prehypophysial portion of sphenoid: absent (0); present (1).
125. Canal for efferent pseudobranchial artery within basicranial cartilage: absent (0); present (1).
126. ***Foramen in basicranium for hypophysis/internal carotids: absent (0); present (1)**

Modified from Mai86 (Chs J22+H17), Did95 (Ch. 51)

This replaces state 2 of ‘internal carotids’ character (see below)

In extant holocephalians the ventral lobe of the pituitary is isolated outside the neurocranium [36]; a hard tissue correlate of this is the absence of a hypophyseal foramen in the basicranium [69]. This foramen is present in some Palaeozoic holocephalians e.g. †*Helodus* [27].

127. **Entrance of internal carotids: through separate openings flanking the hypophyseal opening or recess (0); through a common opening at the central midline of the basicranium (1).**

This character is contingent upon the presence of character 119.

128. **Internal carotids: entering single or paired openings in the basicranium from a posterolateral angle (0); entering basicranial opening(s) head-on from an extreme, lateral angle (1)**

This character is contingent upon the presence of character 119. State 2 (absent) has been removed.

129. Ascending basisphenoid pillar pierced by common internal carotid: absent (0); present (1).
130. Spiracular groove on basicranial surface: absent (0); present (1).
131. Spiracular groove on lateral or transverse wall of jugular canal: absent (0); present (1).
132. Spiracular groove open (0); enclosed by spiracular bar or canal (1).
133. Orbit larger than otic capsule: absent (0); present (1).
134. Postorbital process and arcade: absent (0); present (1).
135. Postorbital process and arcade short and deep - width not more than maximum braincase width (excluding arcade) (0); process and arcade wide - width exceeds maximum width of braincase, and anteroposteriorly narrow (1); process and arcade massive (2); arcade forms postorbital pillar (3).
136. Postorbital process downturned, with anhedral angle relative to basicranium: absent (0); present (1).
137. Jugular canal diameter small (0); large (1); canal absent (2).
138. Canal, likely for trigeminal nerve (V) mandibular ramus, passes through the postorbital process from proximal dorsal entry to distal and ventral exit: absent (0); present (1).
139. Postorbital process articulates with palatoquadrate: absent (0); present (1).
140. Trigemino-facial recess: absent (0); present (1).
141. Jugular canal long, extends throughout most of otic capsule wall posterior to the postorbital process (0); short and/or groove present on exterior of otic wall (1); absent, path of jugular removed from otic wall (2).
142. C-bout notch separates postorbital process from supraotic shelf: absent (0); present (1).
143. Hyoid ramus of facial nerve (N. VII) exits through posterior jugular opening: absent (0); present (1).
144. Periotic process: absent (0); present (1).
145. Relative position of jugular groove and hyomandibular articulation: hyomandibula dorsal or same level (i.e. on bridge) (0); jugular vein passing dorsal or lateral to hyomandibula (1).
146. Transverse otic process: absent (0); present (1).
147. Craniospinal process: absent (0); present (1).
148. Hyomandibula articulates with neurocranium beneath otic shelf: absent (0); present (1).
149. Postotic process: absent (0); present (1)
150. Otic capsule extends posterolaterally relative to occipital arch: absent (0); present (1)
151. Otic capsules: widely separated (0); approaching dorsal midline (1)
152. Otic capsules project anteriorly between postorbital processes: absent (0)	present (1)
153. Endocranial roof anterior to otic capsules domelike, smoothly convex dorsally and anteriorly absent (0); present (1).
154. Roof of skeletal cavity for cerebellum and mesencephalon significantly higher than dorsal-most level of semicircular canals: absent (0); present (1).
155. Roof of the endocranial space for telencephalon and olfactory tracts offset ventrally relative to level of mesencephalon: absent (0); present (1).
156. Labyrinth cavity separated from the main neurocranial cavity by a cartilaginous or ossified capsular wall (0); skeletal medial capsular wall absent (1).
157. External (horizontal) semicircular canal joins the vestibular region dorsal to posterior ampulla (0); joins level with posterior ampulla (1).
158. Angle of external semicircular canal: in lateral view, straight line projected through canal intersects anterior ampulla, external ampullae, and base of foramen magnum: absent (0); present (1)
159. Left and right external semicircular canals approach or meet the posterodorsal midline of the hindbrain roof: absent (0); present (1).
160. Preampullary portion of posterior semicircular canal absent (0); present (1).
161. ***Lagenar chamber distinct from sacculus: absent (0); present(1)**

In many gnathostomes including extant elasmobranchs a distinct lagenar chamber buds off the saccular chamber; this is absent in living holocephalians [70,71]. A lagenar chamber is reported as absent in †*Iniopera* [25]*,* †*Helodus* [61], and †*Kawichthys* [23], and appears to be absent based on figures of †*Ozarcus* (†*‘Cobelodus’* FMNH 13242)[59] and †*Dwykaselachus* [14].

162. Crus commune connecting anterior and posterior semicircular canals: present (0); absent (1).
163. Sinus superior: absent or indistinguishable from union of anterior and posterior canals with saccular chamber (0); present, elongate and nearly vertical (1).
164. Lateral cranial canal: absent (0); present (1).
165. Endolymphatic ducts: posteriodorsally angled tubes (0); tubes oriented vertically	through endolymphatic fossa/posterior dorsal fontanelle (1).
166. Posterior dorsal fontanelle connected to persistent otico-occipital fissure (0); posterior tectum separates fontanelle from fissure (1).
167. Subcircular endolymphatic foramen: absent (0); present (1).
168. External opening for endolymphatic ducts anterior to crus commune: absent (0) present(1).
169. Dorsal otic ridge: absent (0); present (1).
170. Dorsal otic ridge forms a crest posteriorly: absent (0); present (1).
171. Endolymphatic fossa: absent (0); present (1).
172. Endolymphatic fossa elongate (slot-shaped), dividing dorsal otic ridge along midline: absent (0); present (1).
173. Perilymphatic fenestra within the endolymphatic fossa: absent (0); present (1).
174. Ventral cranial fissure: absent (0); present (1).
175. Endoskeletal intracranial joint: absent (0); present (1).
176. Metotic (otic-occipital) fissure: absent (0); present (1).
177. Vestibular fontanelle: absent (0); present (1).
178. Hypotic lamina: absent (0); present (1).
179. Glossopharyngeal nerve path: directed laterally, across floor of the saccular chamber and exits via foramen in side wall of the otic capsule (0); directed posteriorly, and exits through metotic fissure or foramen in posteroventral wall of otic capsule (1); exits laterally through a canal contained ventrally (floored) by the hypotic lamina (2); exits through a foramen anterior to the posterior ampulla (3).
180. Glossopharyngeal and vagus nerves share common exit from neurocranium: absent (0); present (1).
181. Basicranial morphology: platybasic (0); tropibasic (1).
182. Channel for dorsal aorta and/or lateral dorsal aortae passes through basicranium (0): external to basicranium (1).
183. Dorsal aorta divides into lateral dorsal aortae posterior to occipital level (0); anterior to level of the occiput (1).
184. Posterior openings of lateral aortic canals positioned lateral to occipital cotylus: absent (0); present (1).
185. Ventral portion of occipital arch wedged between rear of otic capsules: absent (0); present (1).
186. Dorsal portion of occipital arch wedged between otic capsules: absent (0); present (1).
187. Occipital crest anteroposteriorly elongate, and extends from the roof of the posterior tectum: absent (0); present (1).

*Axial and appendicular skeleton*

188. Calcified vertebral centra: absent (0); present (1).
189. **Chordacentra: absent (0); present (1).**

In all extant holocephalians except for callorhinchids the notochordal sheath is formed from calcified rings (captured in this and the character below). We have scored *Callorhincus* present for this character due to its calcified basidorsal and basiventral cartilages [36], but absent for the next character. Rings are clearly present in some extinct holocephalians (e.g. †*Elasmodus*). Most specimens of †*Ischyodus quenstedti* appear to lack notochordal rings but in one undescribed specimen attributed to the genus ([31], fig. 5) notochordal rings are clearly present; we have scored it uncertain (?). We have scored †*Chondrenchelys* present for this character following the interpretation of [30] and [26]. Following Coates et al. [14] we have scored chordacentra as present in the symmoriiform †*Damocles* and absent in †*Akmonistion*.

190. **Chordacentra polyspondylous and consist of narrow closely packed rings: absent (0); present (1).**

This character refers specifically to the state observed in living and some fossil holocephalians It is contingent on the presence of chordacentra.

191. **Synarcual: absent (0); present (1).**

†*Chimaeropsis* and †*Elasmodectes willetti* were scored as present on the basis of images [31,53].

192. Macromeric dermal pectoral girdle (0); micromeric or lacking dermal skeleton entirely (1).
193. Macromeric pectoral dermal skeleton forms complete ring around the trunk: present	(0); absent (1).
194. Median dorsal plate: absent (0); present (1).
195. Scapular process (dorsal) of shoulder endoskeleton: absent (0); present (1).
196. ***Scapulocoracoids fused ventrally: absent (0); present (1)**

Did95 (Ch. 50)

In extant holocephalians, the left and right scapulocoracoids antimeres are fused ventrally [36]. This is also the case in many extant elasmobranchs (e.*g. [51]),* although in most Palaeozoic chondrichthyans they are separated (e.g. †*Akmonistion* [72]). This appears to be present in †*Chimaeropsis paradoxus* [31,53]*,* but is difficult to judge in may articulated fossil holocephalians due to lateral crushing.

197. Ventral margin of separate scapular ossification: horizontal (0); deeply angled (1).
198. Cross sectional shape of scapular process: flattened or strongly ovate (0); subcircular (1).
199. Scapular process with posterodorsal process. Absent (0); present (1).
200. Coracoid process: absent (0); present (1).
201. Procoracoid mineralisation: absent (0); present (1).
202. Fin base articulation on scapulocoracoid: stenobasal, deeper than wide (0); eurybasal, wider than deep (1).
203. Pectoral fin articulation monobasal (0); dibasal (1); three or more basals (2).
204. ***Anterior pectoral radials fused into block that articulates with propterygium: ansent (0); present (1)**

Did95 (Ch. 64)

This is present in extant holocephalians [36] and in †*Squaloraja* [30]. It is unclear in †*Debeerius* [17] but a similar cartilage appears to be present in †*Helodus* [27,67]. It is absent in †*Chondrenchelys* [26].

205. Metapterygium pectinate subtriangular plate or bar supporting numerous (six or more) radials along distal edge: absent (0); present (1).
206. Metapterygial whip absent (0); present (1).
207. Biserial pectoral fin endoskeleton: absent (0); present (1).
208. Propterygium perforated: absent (0); present (1).
209. Pelvic girdle with fused puboischiadic bar: absent (0); present (1).
210. ***First two to three pelvic fin radials fused into basipterygial process: absent (0); present (1)**

Did95 (Ch. 65)

This unites living chimaeroids (Didier, 1995). It appears to be absent in †*Squaloraja* [30] and other fossil holocephalians where the pelvic fins are preserved.

211. Mixipterygial/mixopterygial claspers: absent (0), present (1).
212. ***Denticles on pelvic claspers: absent (0); present (1)**

Modified from Did95 (Ch. 75)

Chimaerids have denticles on the pelvic claspers, living callorhinchids do not but they are present in †*Squaloraja* [30]. †*Iniopera* and †*Damocles* are scored 1, reflecting the dermal elements identified at the tip of their pelvic claspers [15,60]. Only applicable to taxa with mixopterygial claspers.

213. ***Pelvic claspers multifid: absent (0); present (1)**

Modified from Did95 (Ch. 101)

In *Chimaeridae* the pelvic claspers are either bifid or trifid. This is not the case in *Callorhinchus* [36].

214. **Pre-pelvic clasper or tenaculum: absent (0); present (1).**

This character has been coded as present in †*Squaloraja* based on the enlarged denticles on its pectoral girdle interpreted as a pelvic clasping organ [30].

215. ***Pre-pelvic tenaculum: simple (0); complex (1)**

Extant callorhinchids have complex pre-pelvic tenaculae, in contrast to the simple blade-like tenaculae of chimaerids [36]. This is contingent on having pre-pelvic claspers.

216. Number of dorsal fins, if present: one (0); two (1); one, extending from pectoral to anal fin level (2).
217. Brush complex of bilaterally distributed calcified tubes flanking or embedded in calcified cartilage core: absent (0); present (1).
218. Posterior or pelvic-level dorsal fin with calcified base plate: absent (0); present (1).
219. Posterior dorsal fin with delta-shaped cartilage: absent (0); present (1).
220. Posterior or pelvic-level dorsal fin shape, base approximately as broad as tall and not broader than other median fins (0); base much longer than fin height, substantially longer than other median fins (1).
221. Anal fin: absent (0); present (1).

In living chimaeroids an anal fin is present only in *Callorhincus* and *Neoharriotta* . It is present in †*Isychodus quenstadti* [31,53], which has a *Callorhincus*-like tail morphology. It is unclear whether it is present in either species of †*Elasmodectes* or †*Chimaeropsis* [31,33,34,53].

222. ***Fleshy postanal tail pad: absent (0); present (1)**

Did95 (Ch. 102)

Synapomorphy of *Chimaera* and *Hydrolagus* [36].

223. Caudal radials restricted to axial lobe (0); extend beyond level of body wall and deep into hypochordal lobe (1).
224. Caudal neural and/or supraneural spines or radials short (0); long, expanded, and supporting high aspect-ratio (lunate) tail with notochord extending to posterodorsal extremity (1); notochord terminates pre-caudal extremity, neural and heamal radial lengths near symmetrical and support epichordal and hypochordal lobes respectively (2).
225. ***Caudal fin shape: heterocercal (0); homocercal (1)**

Did95 (Ch. 81)

Alone among extant holocephalians *Callorhincus* has a heterocercal tail. In extinct taxa this is also present in †*Debeerius* and in †*Ischyodus* [17,31,53]. Extant chimaerids have a homocercal tail: seemingly this is shared by †*Elasmodectes* [33,53].

226. *** Tubercles on supracaudal lobe of the tail in males: absent (0); present (1)**

Did95 (Ch. 96)

This is a synapomorphy of *Rhinochimaera.*

*Spines: fins, cranial and elsewhere*

227. Dorsal fin spine or spines: absent (0); present (1).
228. Dorsal fin spine at anterior (pectoral level) location only: absent (0); present (1).
229. Dorsal fin spine apex curved posteriorly: absent (0); present (1).
230. Anterior dorsal fin spine leading edge concave in lateral view: absent (0); present (1).
231. ***Trabecular dentine in lateral walls of dorsal fin spine: reduced or absent (0); present (1)**

Did95 (Ch. 49)

Extant holocephalians have a reduced layer of trabecular dentine in the lateral walls of their dorsal fin spine [36], with it being limited to the anterior keel in *Callorhincus* [73]. This character is scored inapplicable for taxa without dorsal fin spines.

232. Anal fin spine: absent (0); present (1).
233. Pectoral fin spines: absent (0); present (1).
234. Median fin spine insertion: shallow, not greatly deeper than dermal bones/ scales (0); deep (1).
235. Fin spines with ridges: absent (0); present (1).
236. Fin spines (dorsal) with rows of large denticles: absent (0); on posterior surface (1); on lateral surface (2).

In myriacanthids the lateral sides of the fin spine are covered in tubercles, coded here as character state 2 [30]. In †*Acanthorhina* this is unclear due to preparation [65]. it is scored unknown ?.

**Supplementary References**.

1.	Zhu M *et al.* 2013 A Silurian placoderm with osteichthyan-like marginal jaw bones. *Nature* **502**, 188–193. (doi:10.1038/nature12617)

2.	Gradstein FM, Ogg JG, Schmitz M, Ogg G. 2021 *The Geologic Time Scale 2020*. Elsevier Science.

28.	Pradel A, Denton JSS, Janvier P, Maisey JG, editors. 2021 *Ancient fishes and their living relatives: a tribute to John G. Maisey*. München: Verlag Dr. Friedrich Pfeil.

29.	Duffin CJ, Garassino A, Pasini G. 2023 *Squaloraja* Riley 1833 (Holocephala: Squalorajidae) from the Lower Jurassic of Osteno Konservat-Lagerstätte (Como, NW Italy). *Nat. Hist. Sci.* **10**. (doi:10.4081/nhs.2023.642)

30.	Patterson C. 1965 The Phylogeny of the Chimaeroids. *Philos. Trans. R. Soc. Lond. B. Biol. Sci.* **249**, 101–219.

31.	Villalobos-Segura E, Stumpf S, Türtscher J, Jambura PL, Begat A, López-Romero FA, Fischer J, Kriwet J. 2023 A Synoptic Review of the Cartilaginous Fishes (Chondrichthyes: Holocephali, Elasmobranchii) from the Upper Jurassic Konservat-Lagerstätten of Southern Germany: Taxonomy, Diversity, and Faunal Relationships. *Diversity* **15**, 386. (doi:10.3390/d15030386)

32.	Fraas E. 1910 Chimäridenreste aus dem oberen Lias von Holzmaden. *Jahresh. Ver. Für Vaterl. Naturkunde Württ.* **66**, 55–63.

33.	Lauer B, Lauer R, Bernard E, Duffin C, Popov E, Ward D. 2019 *Observations on the Mesozoic chimaeroid, Elasmodectes Newton 1878*.

34.	Ward D, Richter M, Popov E, Bernard E. 2014 *The first holomorphic fossil chimaeroid fish (Chondrichthyes, Holocephali) from Africa.*

35.	Woodward AS. 2014 *The Fossil Fishes of the English Chalk*. Cambridge: Cambridge University Press. (doi:10.1017/CBO9781139680868)

36.	Didier DA. 1995 Phylogenetic systematics of extant chimaeroid fishes (Holocephali, Chimaeroidei). American Museum novitates ; no. 3119.

47.	1973 *Interrelationships of Fishes: Greenwood, R.S. Miles, Colin Patterson*. Linnean Society of London.

53.	Duffin C, Ward D, Lauer B, Lauer R. 2020 *The Holocephalian faunas of the Late Jurassic lithographic limestones of SW Germany*.

54.	Dean B. 1909 Studies on fossil fishes (sharks, chimaeroids and arthrodires. Memoirs of the AMNH ; v. 9, pt. 5.

55.	Dean B. 1894 Contributions to the morphology of cladoselache (Cladodus). *J. Morphol.* **9**, 87–114. (doi:10.1002/jmor.1050090103)

56.	Tomita T. 2015 Pectoral Fin of the Paleozoic Shark, Cladoselache: New Reconstruction Based on a Near-Complete Specimen. *J. Vertebr. Paleontol.* **35**, 1–4.

57.	Woodward AS, White EI. 1938 XLIII.—The dermal tubercles of the Upper Devonian shark, Cladoselache. *Ann. Mag. Nat. Hist.* **2**, 367–368. (doi:10.1080/00222933808526863)

58.	Harris JE. 1938 *The Dorsal Spine of Cladoselache: The Neurocranium and Jaws of Cladselache*.

59.	Maisey JG. 2007 THE BRAINCASE IN PALEOZOIC SYMMORIIFORM AND CLADOSELACHIAN SHARKS. *Bull. Am. Mus. Nat. Hist.* **2007**, 1–122. (doi:10.1206/0003-0090(2007)307[1:TBIPSA]2.0.CO;2)

60.	Zangerl R, Case G. 1976 Cobelodus aculeatus (Cope), an anacanthous shark from Pennsylvanian black shales of North America. *Palaeontogr. A* **154**, 107–157.

61.	Pradel A, Denton JSS, Janvier P, Maisey JG, editors. 2021 *Ancient fishes and their living relatives: a tribute to John G. Maisey*. München: Verlag Dr. Friedrich Pfeil.

67.	Stahl BJ. 1999 *Chondrichthyes III: Holocephali*. Pfeil.

68.	PATTERSON C. 1992 Interpretation of the toothplates of chimaeroid fishes. *Zool. J. Linn. Soc.* **106**, 33–61. (doi:10.1111/j.1096-3642.1992.tb01239.x)

69.	Maisey JG. 1986 Heads and Tails: A Chordate Phylogeny. *Cladistics* **2**, 201–256. (doi:10.1111/j.1096-0031.1986.tb00462.x)

70.	Maisey JG, Lane JA. 2010 Labyrinth morphology and the evolution of low-frequency phonoreception in elasmobranchs. *Comptes Rendus Palevol* **9**, 289–309. (doi:10.1016/j.crpv.2010.07.021)

71.	Maisey JG. 2001 A primitive chondrichthyan braincase from the Middle Devonian of Bolivia. In *Major Events in Early Vertebrate Evolution*, CRC Press.

72.	Coates MI, Sequeira SEK. 2001 A new stethacanthid chondrichthyan from the lower Carboniferous of Bearsden, Scotland. *J. Vertebr. Paleontol.* **21**, 438–459. (doi:10.1671/0272-4634(2001)021[0438:ANSCFT]2.0.CO;2)

73.	Jerve A, Johanson Z, Ahlberg P, Boisvert C. 2014 Embryonic development of fin spines in Callorhinchus milii (Holocephali); implications for chondrichthyan fin spine evolution. *Evol. Dev.* **16**, 339–353. (doi:10.1111/ede.12104)

**Supplementary Figure Captions.**

**Figure S1. Parsimony Analysis Results on Morphological Dataset.** Figure shows the Adams consensus and maximum agreement subtree resulting from the analysis of the morphological dataset in PAUP. Daggers denote extinct taxa.

**Figure S2. Bayesian Analysis Results on Morphological Dataset.** Figure shows the 50% majority rule consensus tree resulting from the analysis of the morphological dataset in MrBayes. Daggers denote extinct taxa. PS=Posterior support.

**Figure S3. Maximum Likelihood Analysis of the Molecular Dataset.** Figure shows the phylogeny of living holocephalians estimated in analysis of the molecular dataset under a maximum likelihood criterion in IQ-TREE2. BS=Ultrafast bootstrap support. Asterisks denote type species of genera recognized in this study.
