## Supplementary figures and images for "The Paleozoic assembly of the holocephalian body plan far preceded post-Cretaceous radiations into the ocean depths"

### Supplementary Figure 1

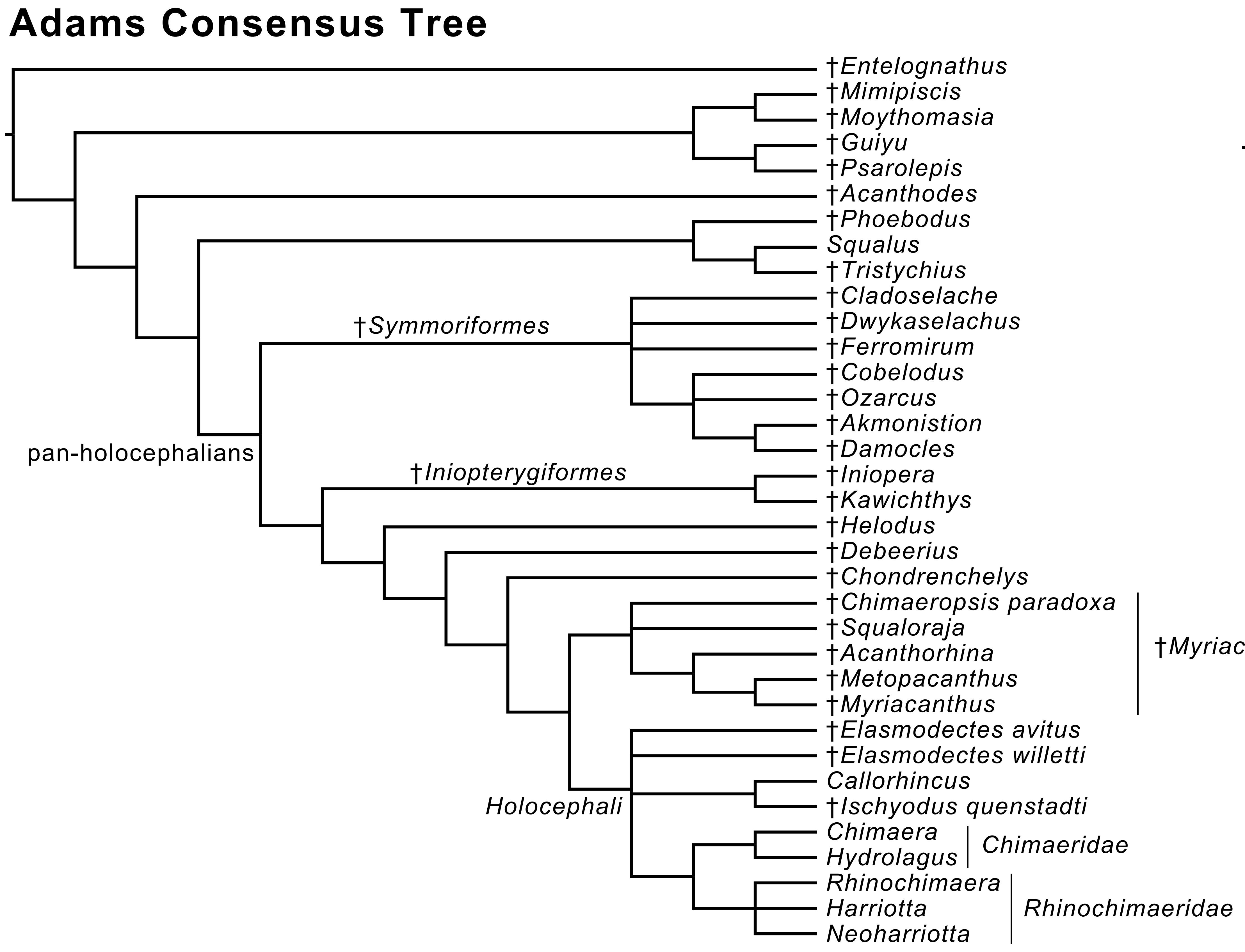

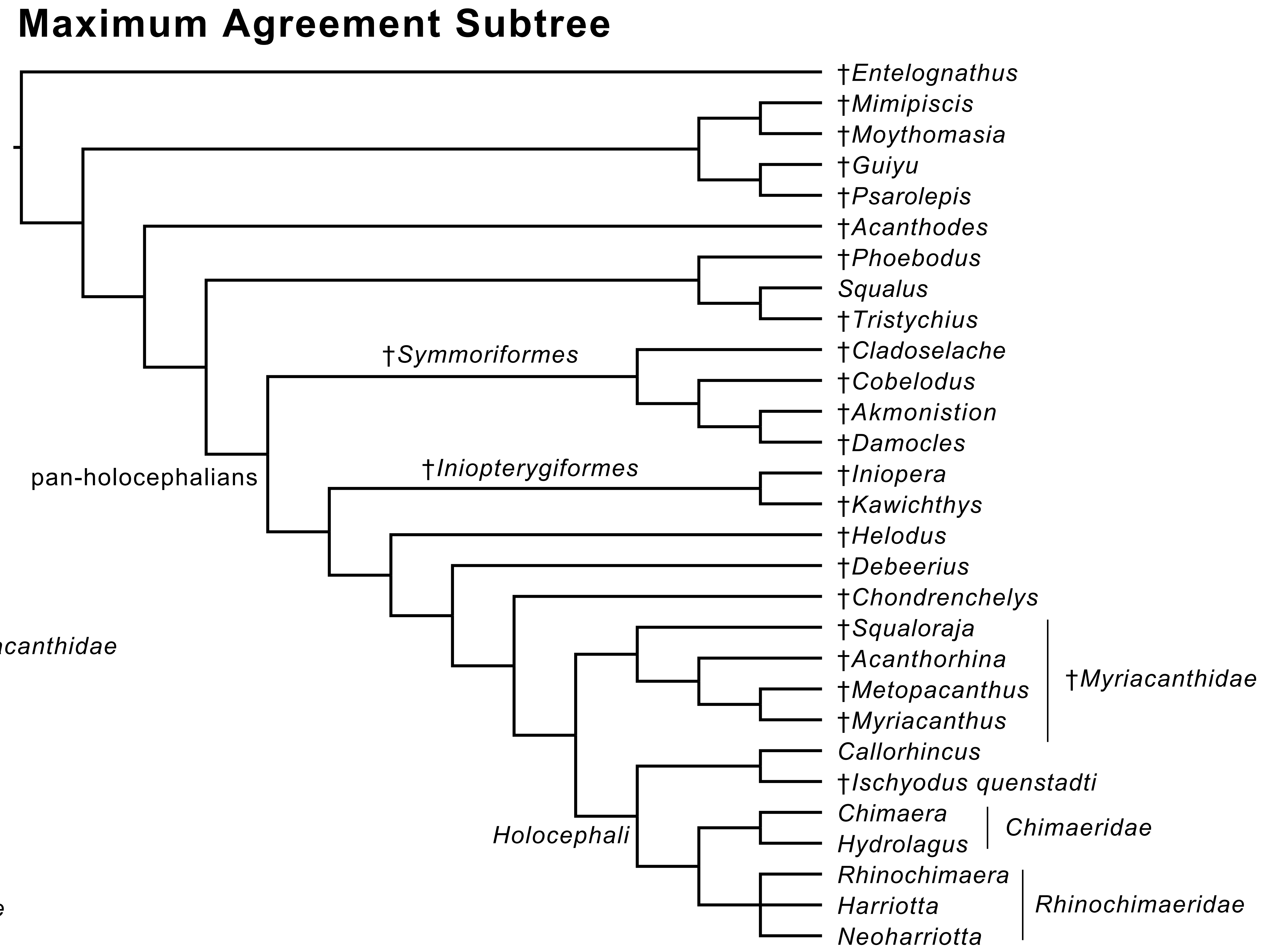

### Supplementary Figure 2

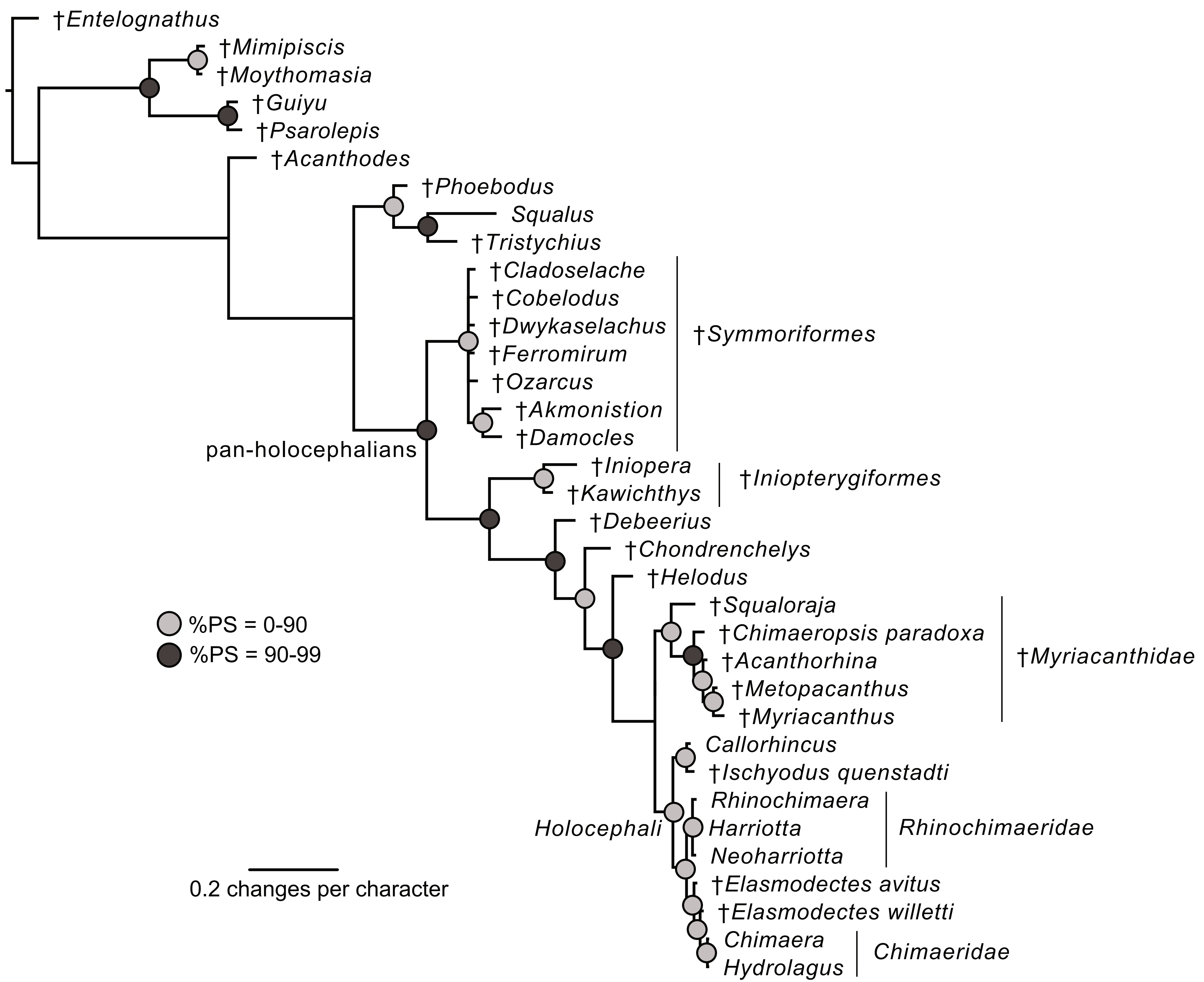

### Supplementary Figure 3

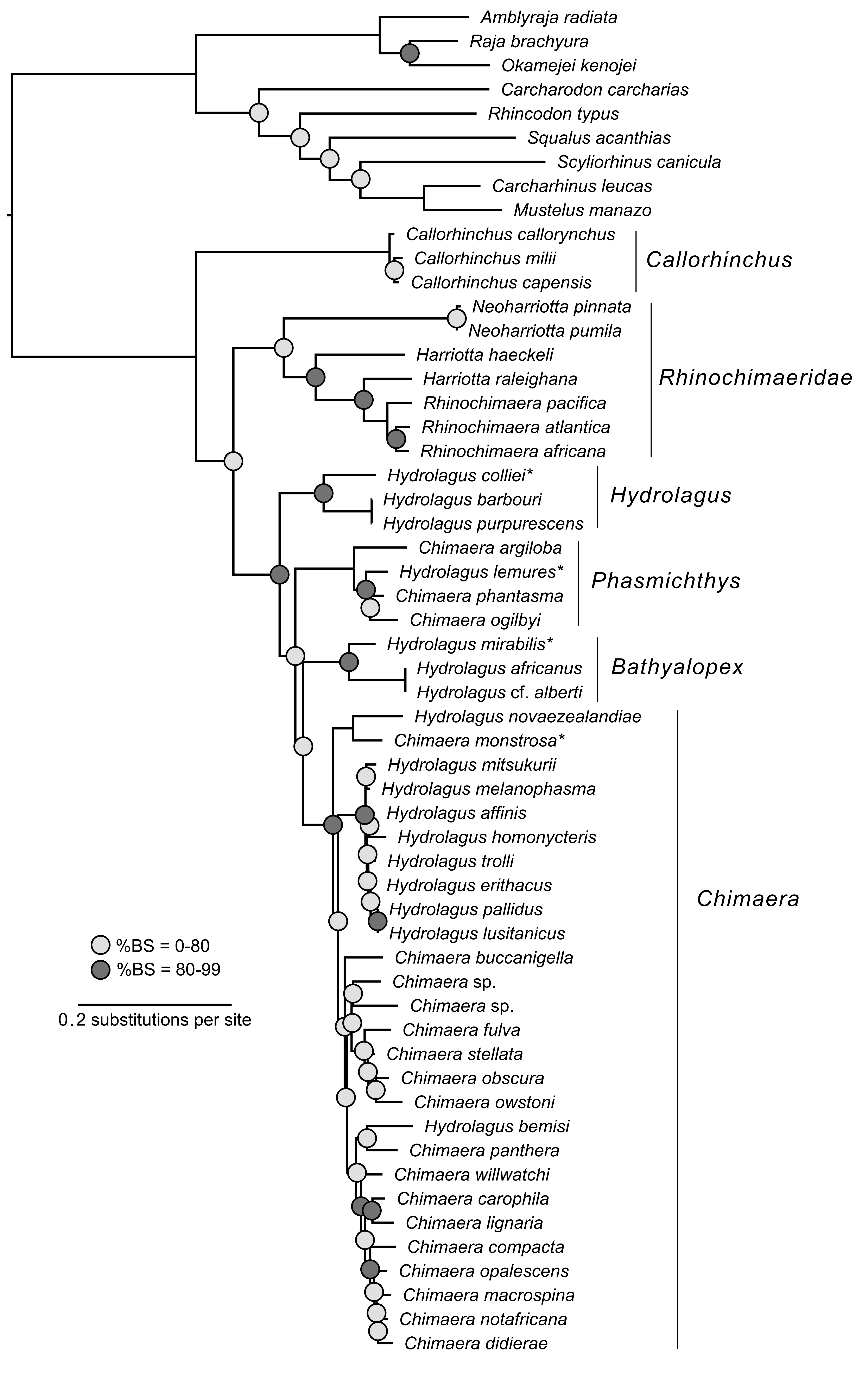
